# Pareto-optimal synthesis of multiple glycans in Golgi compartments

**DOI:** 10.1101/2025.03.14.643235

**Authors:** Aashish Satyajith, Mukund Thattai

## Abstract

Tree-shaped sugar chains called glycans are covalently attached to proteins in the plasma membrane of eukaryotic cells. They engage in multiple functions such as adhesion, signaling, etc. at the cell surface, so their correct manufacture is of vital importance. Glycans are assembled step by step through enzyme-catalyzed monomer addition reactions as they transit through compartments of the Golgi apparatus. The constraint of fixed residence times across Golgi compartments creates tradeoffs: a short residence time does not give complex glycans enough time for reactions to run to completion, while a long residence time might execute undesirable reactions, which inevitably occur due to enzyme promiscuity. It is not clear how the Golgi reconciles this trade-off.

To study this, we devise a from-first-principles model of glycan manufacture with a focus on maximizing glycan yield. Pareto-optimal solutions are a class of solutions that reconcile trade-offs in simultaneously maximizing the yield of multiple glycans. We explore Pareto-optimal solutions for multiple glycan manufacture, and show that small changes in residence times or relative glycan yields can change the optimal enzyme distribution. Together, these results establish Pareto optimality as a unifying framework for interpreting Golgi compartment organization, and for controlling glycan outputs in industrial settings such as manufacture of biotherapeutics, many of which are glycosylated.

## Introduction

Proteins and lipids across all domains of life, particularly those on the cell surface, have branched sugar polymers known as glycans covalently attached to them [1–3]. These structures play important roles in cellular and organismal biology [4]. For example, glycans influence the biochemical and biophysical properties of cell surfaces, mediate cell-cell interactions in tissues [3], and underlie host-pathogen specificity [5]. Incorrect glycan formation has also been linked to many non-pathogenic diseases in humans [6–8].

In eukaryotic cells, proteins destined for the plasma membrane enter secretory pathway at the endoplasmic reticulum (ER), and then move through the Golgi apparatus [9]. As they pass through the Golgi, secreted proteins are subjected to chemical modifications called glycosylation [10]: the covalent addition of sugar monomers to construct tree-like polymeric structures known as glycans. These additions are catalyzed by enzymes called glycosyltransferases [10], which are localized in the lumen of the ER and Golgi. The Golgi itself is organized as an ordered series of compartments, with each compartment containing distinct sets of glycosyltransferase enzymes [11, 12]. This spatially ordered architecture effectively turns the Golgi into a biochemical assembly line, with successive cisternae acting as distinct batch reactors. Unlike template-directed processes such as DNA replication or translation, glycan synthesis lacks a strict molecular blueprint, so control must be exerted through enzyme localization, reaction kinetics, and transport times.

The Golgi manufactures a vast repertoire of glycans at a given time [13]. In the maturation model of the Golgi apparatus, glycan substrates undergoing glycosylation in the same cisterna are exposed to its environment for the same amount of time before the cisterna matures [14]. The amount of time a glycan resides in a compartment is called its residence time. A short residence time might not suffice for the manufacture of a complex glycan to run to completion. A long residence time might be detrimental for fully grown glycans, since promiscuous enzymes might catalyze undesirable reactions [15]. Both these events have varying functional implications, such as reducing the half-life of glycoprotein in circulation [16, 17], altering immune response [18], etc. The protein quality control mechanisms in the Golgi predominantly retrieve or degrade misfolded, mislocalized, or “orphaned” proteins that reach the Golgi [19]. The mechanisms are not able to detect and correct inadequately processed or overprocessed glycans as described above. Both these events result in reduced yield of desired glycans, and represent significant biosynthetic costs to the cell [20].

In addition to compartment residence times, the enzyme distribution across Golgi also influences glycan yield. Previous work from our group rigorously demonstrated that enzyme compartmentalization leads to greater control of glycan assembly [21]. Systems models of glycosylation such as [13, 22–26] have physical parameters (residence times, number of compartments) and chemical parameters (enzyme rates, distribution across compartments) to understand how both these factors influence the distribution of cell-surface glycans. The parameters are then fit to observed or target distributions [27]. However, the fact that glycan manufacture incurs significant biosynthetic cost to the cell, and the resulting incentive to maximize glycan yield, remains understudied.

Here we explore how the enzyme distribution across the Golgi compartments might maximize glycan yield when there are multiple desired glycans. It might be possible to improve the yield of one glycan at the cost of another. The constraint to maximize glycan yield limits the space of allowed configurations of residence time and enzyme distribution: for a given configuration, no other configuration should be able to generate more of one glycan without decreasing the yields of other glycans. The space of such configurations is called the Pareto set of configurations [28]. The notion of Pareto optimality has wide-ranging applications in engineering [29], social sciences [30], and several areas of biology such as ecological studies [31–33], neuroscience [34] etc.

We consider an abstract formulation of glycan manufacture, and devise a simple chemical kinetic model for the synthesis of a glycan. This allows us to infer the optimal choice of enzyme distribution and residence times for that glycan. The requirement for multiple glycans to be manufactured – each of which can have differing residence time requirements for their synthesis – brings in trade-offs. Pareto-optimal solutions are a class of solutions that reconcile this trade-off. The Pareto set reveals that small shifts in compartment residence times or relative yields of glycans changes the optimal enzyme distribution. Studying the Pareto set has practical applications: in the pharmaceutical industry there is value in controlling glycosylation to achieve better antibody efficacy, selective targeting of effector cells, etc. [35–37]. More broadly, our work connects glycan biosynthesis to a growing body of biophysical studies that aim to understand general design principles in multi-objective cellular processes.

## Materials AND Methods

### Model development

We adopt the cisternal maturation model of the Golgi apparatus [14, 38]: the residence time in a compartment is the same for all glycoproteins that transit that compartment, although the residence times of different compartments can be different. Glycans are grown by requisite (desirable) monomer addition reactions on the structure that contains the root. We assume a constant supply of donor sugar nucleotides. Each glycan has a set of enzymes needed for its manufacture, and we assume that enzymes of one glycan do not catalyze the reactions needed for another glycan. We assume the glycosylation enzymes are not saturated, and therefore the monomer addition reactions obey first-order kinetics [39]. We set the time unit so that the “effective” rate constant of all desirable reactions is 1 unit. The enzymes catalyzing the desirable reactions are promiscuous; they can catalyze reactions other than their primary reaction, albeit at a much smaller rate constant [15]. Let this smaller rate constant be *q*. Any of the enzymes in a compartment can execute undesirable reactions: if there are | *Z*_*j*_ | number of enzymes in compartment *j* that manufacture a glycan, the effective rate constant of undesirable reactions is *q* · | *Z*_*j*_ |.

The amount of each glycan produced is controlled by the enzyme distribution across compartments and the residence times of each compartment.

#### Enzyme splits

Let ℰ_*B*_ be the set of enzymes that are required to synthesize a glycan tree *B*. For an *N* -compartment Golgi apparatus, a split denotes the enzymes present in each of the compartments. Formally, a split 𝒮 is an element of 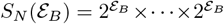, product taken *N* times, where 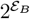 denotes the power set of ℰ_*B*_. Thus the enzyme split of a glycan can be represented as (*Z*_1_, …, *Z*_*n*_) where *Z*_*j*_ denotes the set of enzymes in compartment *j*. We consider only partitions of *E*_*B*_ as splits (i.e. no enzyme appears in more than one compartment), as an approximation of the fact that many Golgi enzymes are localized to one or two Golgi cisternae [40]. An example of enzyme splits is given in Fig. 1A.

**Fig. 1.**
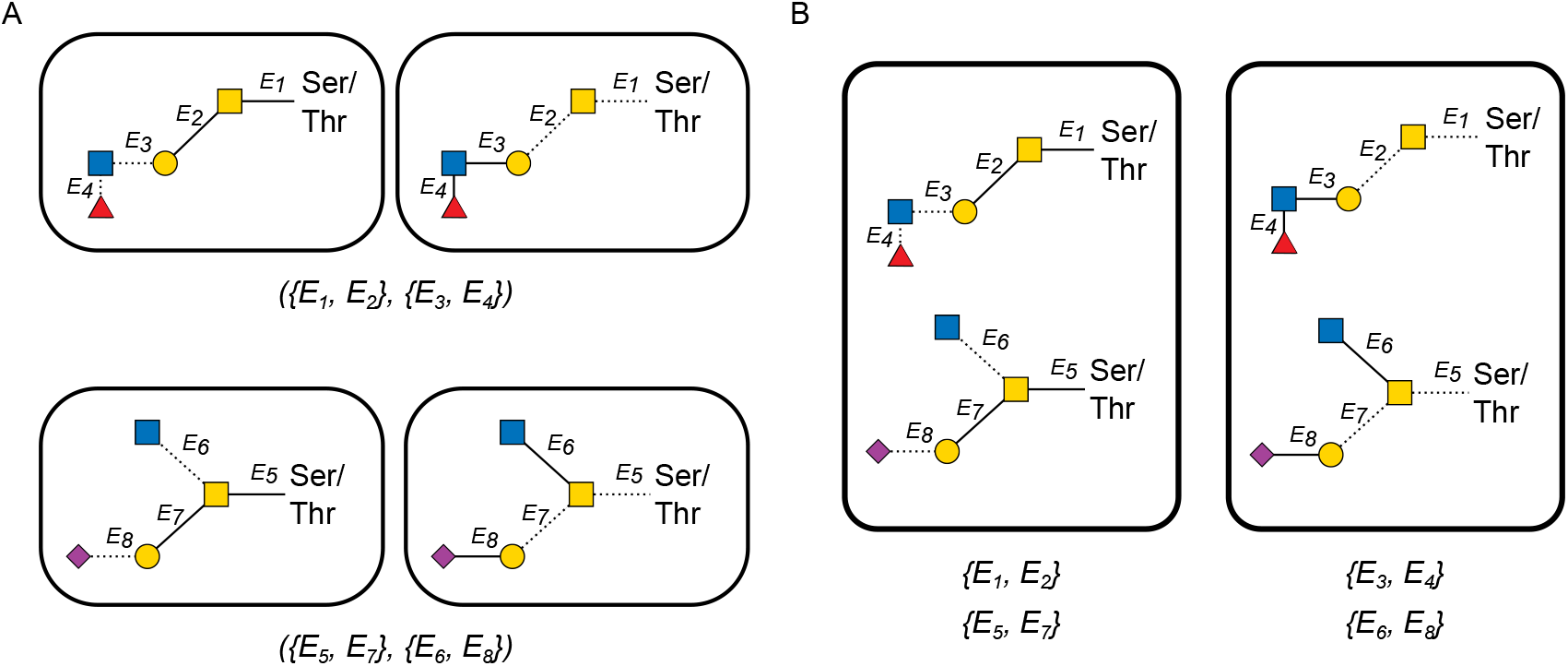
**(A)** Splits for manufacturing glycans in the left and right panels of Figure 2A (top and bottom boxes respectively). The top two boxes denote a Golgi configuration with enzymes *E*_1_ and *E*_2_ in the first compartment, and *E*_3_ and *E*_4_ in the second compartment. The bottom boxes denote a Golgi configuration with enzymes *E*_5_ and *E*_7_ in the first compartment, and *E*_6_ and *E*_8_ in the next compartment. The thick lines in a structure inside a compartment denote reactions that are to happen in that compartment, while dotted lines denote reactions that happen elsewhere. Splits can be represented pictorially or via text, as shown. **(B)** Golgi configurations needed to simultaneously produce the different glycans shown in 2A. In the second compartment, two serial and two parallel reactions need to happen for the correct manufacture of the respective glycans. The reactions will have different optimal residence times, creating trade-offs.

#### Residence times

For an *N* -compartment Golgi apparatus, residence times are specified by a tuple in 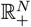.

A *strategy* denotes the splits and the residence times assigned to each compartment. They are represented as a two-tuple (𝒮 𝒯,) where 𝒮 and 𝒯 are elements of *S*_*N*_ (ℰ _*B*_) and 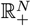 respectively. Once the splits and residence times have been fixed, we can calculate the chance that a given glycan is correctly formed. We call this the *yield* of that glycan.

#### Calculating glycan yield

We assume that the major contributors that decrease yield of a target glycan are required reactions that do not run to completion, and undesirable reactions executed by promiscuous enzymes [39]. We ignore other types of reactions in this work [21]. Since monomer addition reactions are assumed to follow first-order kinetics, the waiting time for reactions follows an exponential distribution. For the complete development of a branch, a monomer addition reaction has to first take place, initiating the creation of that branch, followed by all the other reactions needed to complete that branch. We use this observation to calculate yields in an inductive fashion. Let *a*_1_, …, *a*_*K*_ be the nodes attached to the root node, in a tree *B* that is being grown in a compartment. Denote the subtree rooted at *a*_1_, …, *a*_*K*_ by 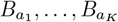 respectively. Suppose the *i*-th branch is to be made in time *t*. Let *t*_1_ be the time at which the *a*_*i*_ monomer addition to the root takes place, starting the growth of that branch. This time can be anywhere from 0 to *t*, i.e. 0 ≤ *t*_1_ ≤ *t*. The rest of the branch has time *t* − *t*_1_ to develop. Thus the *yield of formation* of that branch can be written recursively as

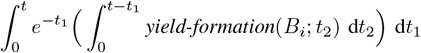

Independent components of a glycan tree can grow at the same time in a compartment, i.e. reactions can proceed in parallel. The parallel growth of the glycan to the right in Fig. 2A is shown in Fig. 2B. The yield of formation of the entire glycan *B* is then given by the product of yields of formation of independent components:

**Fig. 2.**
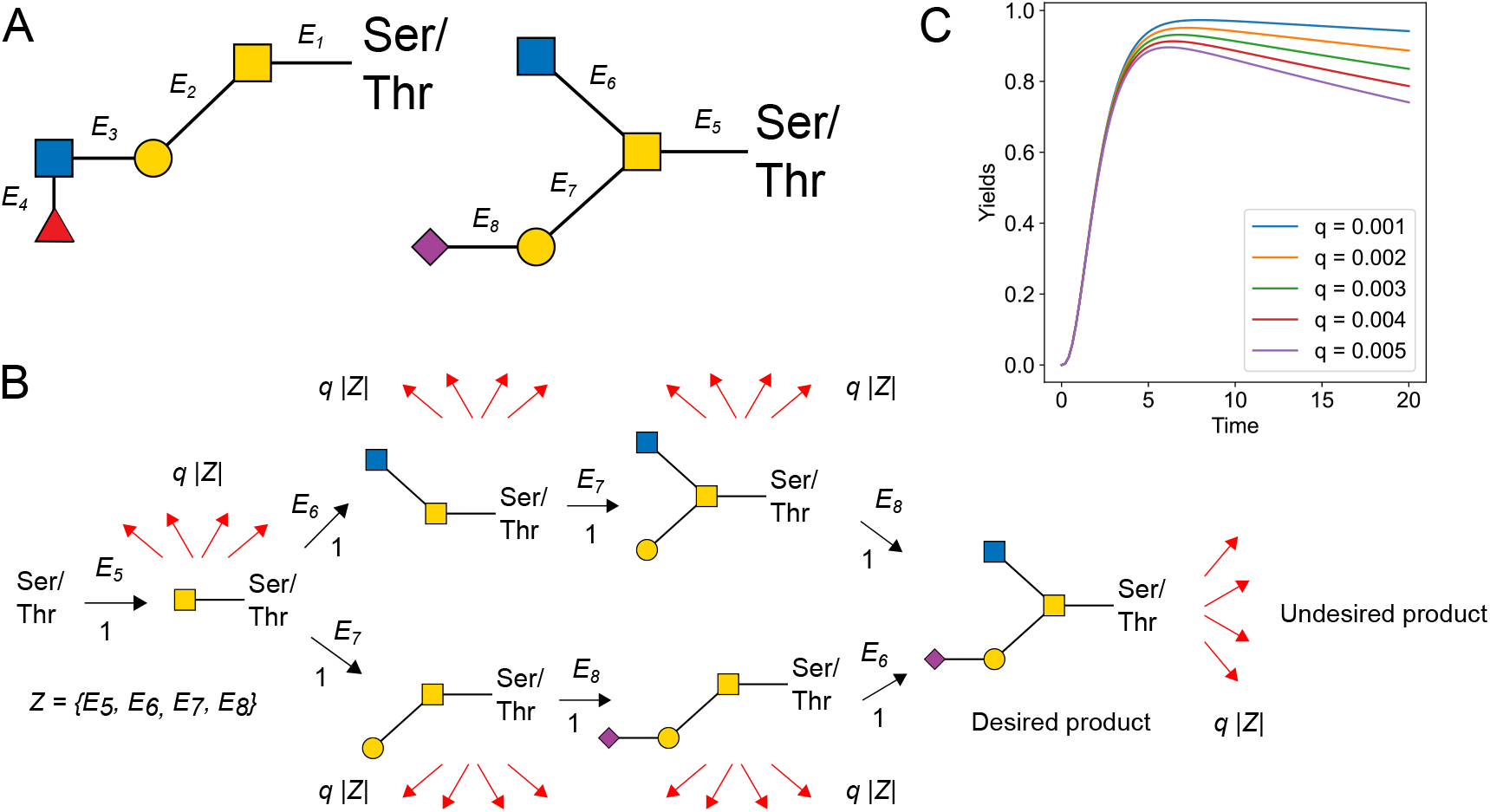
Enzymes catalyze desirable and undesirable reactions, at different rates. **(A)** Examples of glycan structures manufactured on a Ser/Thr residue on the substrate protein. The colored shapes denote sugar monomers [41]. The labels on the edges denote enzymes that are required to catalyze the linkage reaction. The glycan to the left needs four serial reactions, i.e. reactions occurring one after another. In case of the glycan to the right, the reaction catalyzed by *E*_6_ can proceed independently of the other branch, once the *E*_5_ reaction has taken place. **(B)** Different growth orders that can give rise to the glycan to the right of panel A. The numbers beside the arrows indicate the rate constants of reactions. The red arrows denote undesirable reactions. *Z* denotes the set of enzymes needed to manufacture the glycan. **(C)** Yield of the glycan to the right of (A) at different values of the undesirable reaction rate *q* and varying residence times.

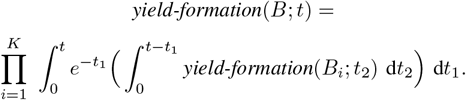

As an example, consider the subtree rooted at the GalNAc monomer (yellow square) in Fig. 2A, right side. The yield of formation of required reactions can be written down as

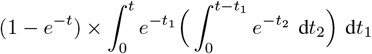

The term 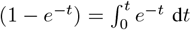 on the left corresponds to the yield of the branch with one reaction. The term on the right corresponds to the yield of two serial reactions. We note that for trees that have only serial reactions or parallel reactions, simpler expressions for yield can exist.

Undesirable reactions occur with rate constant *q*. Since the effective rate constant of undesirable reactions in a compartment with | *Z*_*j*_ | enzymes is *q* · | *Z*_*j*_ |, the chance of an undesirable reaction is 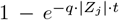. Therefore the chance of it not occurring is 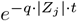.

Reactions are irreversible [21], so the correct formation of a glycan depends on desirable reactions occurring and undesirable reactions not occurring. We assume that desirable or undesirable reactions occur independently, and hence the yield of a glycan is given by the product of these two events. The yield of a glycan *B* under enzyme splits (*Z*_1_, …, *Z*_*N*_) and residence times (*t*_1_, …, *t*_*N*_) is 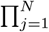 *yield-formation* 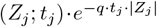.

Define the yield of a strategy (𝒮, 𝒯) as

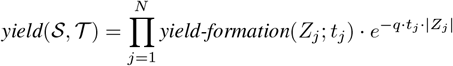

where ( 𝒮 𝒯,) = ((*Z*_1_, …, *Z*_*N*_), (*t*_1_, …, *t*_*N*_)). Hence the objective function for the optimal yield of a glycan *B* can be formulated as:

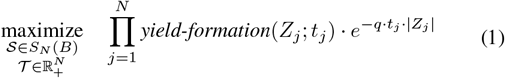

For a fixed split, the yield functions can be shown to be a pointwise log-concave function of residence times (Appendix A). Log-concave functions have a maximum value, and several fast algorithms exist to find the maximizer [42]. Thus, the optimal residence times and enzyme distribution to manufacture a glycan can be found by maximizing the log-yield function corresponding to each split, and picking the split that gives the best yield.

A glycoprotein population can include many glycoforms with varying functional activity [10], all of which are produced by the same Golgi apparatus. Consider a Golgi apparatus producing *K* glycan trees *B*_1_, …, *B*_*K*_. Let 𝒮^1^, …, 𝒮^*K*^ denote the splits of the *K* glycans. We will write the collection of splits 𝒮^1^, …, 𝒮^*K*^ as **𝒮**, in bold type, and write **𝒮**^*i*^ to denote the split corresponding to the *i*^th^ glycan. Once we fix the splits for each of the *K* glycans and the residence times of each compartment, denoted ( 𝒮 𝒯,), we can calculate the yields *y*_1_, …, *y*_*K*_ for each of the glycans.

This process of synthesising multiple glycans can create trade-offs, arising from the fact that the residence times are fixed for each compartment. Fig. 1B gives an example of such a scenario. The optimal residence time for two serial reactions is greater than the optimal residence time for two parallel reactions. Fixing the residence time to be the optimal residence time of one glycan will be detrimental to the other glycan.

The cell will have its own fitness function *f* (*y*_1_, …, *y*_*K*_) based on the yields *y*_1_, …, *y*_*K*_ of the *K* glycans, which we assume is what the cell is optimized for. We assume that if more of a glycan can be manufactured without decreasing the amounts of other glycans, that is advantageous for the cell; in other words, *f* (*y*_1_, …, *y*_*K*_) is a monotonically increasing function of *y*_*i*_ ∀ *i*, keeping *y*_1_, …, *y*_*i*−1_, *y*_*i*+1_, …, *y*_*K*_ fixed. This implies that the Golgi configuration must be from the Pareto set: the set of strategies where no yield can be increased without lowering another. With this assumption, we can define Pareto dominance of strategies.

### Pareto optimality

A strategy is said to *Pareto dominate* another strategy if all the glycans that are being synthesized have either same or better yields using the former strategy than the latter strategy [28]. Formally, let (*u*_1_, …, *u*_*K*_) and (*v*_1_, …, *v*_*K*_) be the yield vectors corresponding to strategies ( **𝒮 𝒯**,) and ( **𝒮**^***′***^, **𝒯** ^′^) respectively. ( **𝒮**^***′***^, **𝒯** ^′^) Pareto dominates ( **𝒮 𝒯**,), written (**𝒮 𝒯**,) ≺ ( **𝒮**^***′***^ **𝒯**, ^′^), if *u*_*i*_ ≤ *v*_*i*_, ∀*i* ∈ [*K*] and ∃*i* | *u*_*i*_ *< v*_*i*_. The set of all strategies that cannot be Pareto dominated is called the Pareto set, and the set of their corresponding yields is called the Pareto front. Given our assumption earlier that *f* (.) is monotonically increasing, we only need to consider the set of Pareto-optimal solutions for further study. Pareto-optimal solutions for multiple glycan manufacture can be obtained by enumerating combinations of splits of the glycans and maximizing convex weightings of the log-yield functions (see Appendix B).

## RESULTS

We first characterize optimal Golgi configurations for a single glycan, starting from a one-compartment system and then extending to a two-compartment Golgi. The two-compartment case is the smallest architecture that exhibits nontrivial tradeoffs when multiple glycans are to be manufactured; unless stated otherwise, all subsequent results use this two-compartment setting.

For a single glycan, the optimal Golgi configuration reduces to that glycan’s optimal split and residence times. Consider the glycan to the right of Fig. 2A being grown in a single compartment. Fig. 2C shows its yield for different residence times and at different values of undesirable reaction rate constant *q*. There is an optimal residence time for its synthesis, and other residence times lead to suboptimal yields. A global maximum is guaranteed the yield function is log-concave in residence times (Appendix A).

This result extends to two compartments. Consider the manufacture of a glycan that needs four serial reactions for its correct synthesis. Fig. 3A plots the yield contours under a 1-3 split and a 2-2 split with respect to residence times in the two compartments. For both the splits, there are optimal residence times in the first and second compartments. The yields of each split at different *q* is plotted in Fig. 3B. The symmetric split achieves the highest yield among all the splits, and demonstrates that some splits achieve better yield than others.

**Fig. 3.**
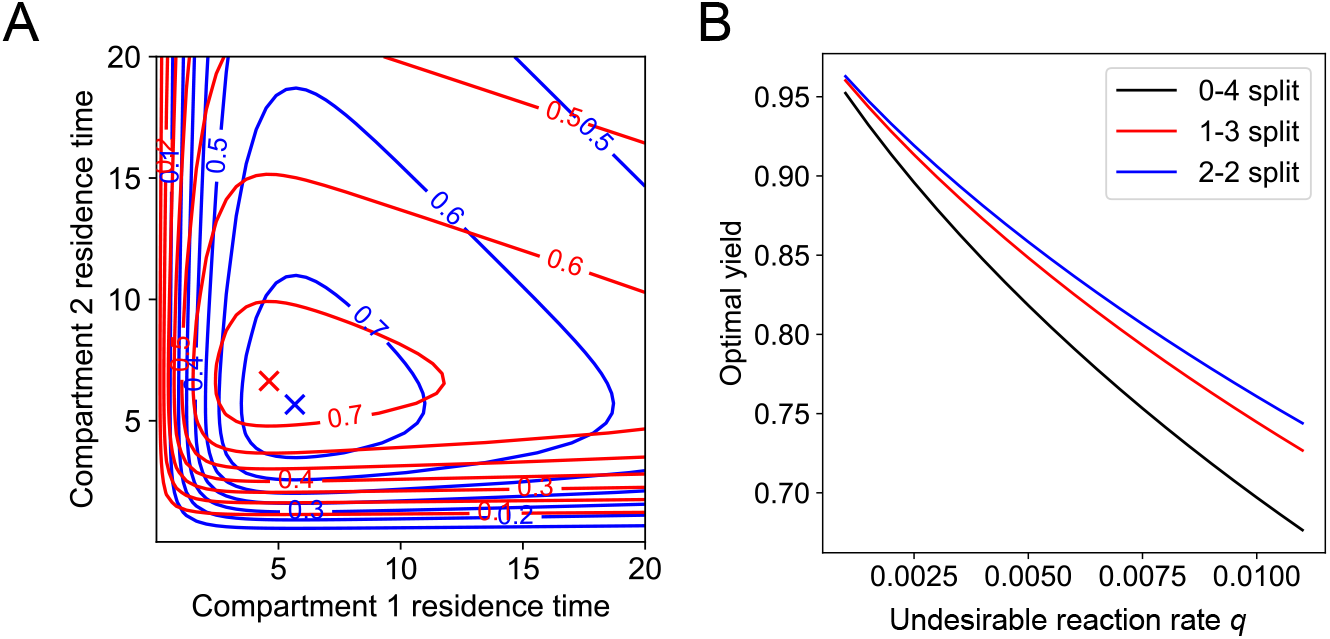
Optimal splits and residence times for a single glycan needing four serial reactions. **(A)** Yields at various residence times for different splits of four serial reactions, at *q* = 0.01. The blue lines denote contours for a 2-2 split (first two reactions in the first compartment, next two in the second compartment) while the red lines denote contours for a 1-3 split (first reaction in the first compartment, three remaining serial reactions in the next compartment). The ‘x’ marks in the same colours denote the optimal residence times of the splits in first and second compartments. The 3-1 split and the 4-0 split need not be considered in the single glycan manufacture case, due to symmetry. **(B)** Optimal yield vs *q* curve for all possible splits of four reactions across two compartments.

We next ask how the optima interact when multiple glycans are manufactured in the same Golgi. To explore this, we chose three glycans with qualitatively different structures, each requiring six reactions after attachment of the root with the glycoprotein substrate (Fig. 4A). The first glycan consists of edges that offer opportunities for parallel growth. The second glycan consists of only serial reactions. The third glycan has three branches which need one, two, and three serial reactions. The optimal residence times for the Pareto-optimal splits are shown as thick dots in Fig. 4D, and illustrates the trade-off: different glycans have different optimal residence times across the two compartments.

**Fig. 4.**
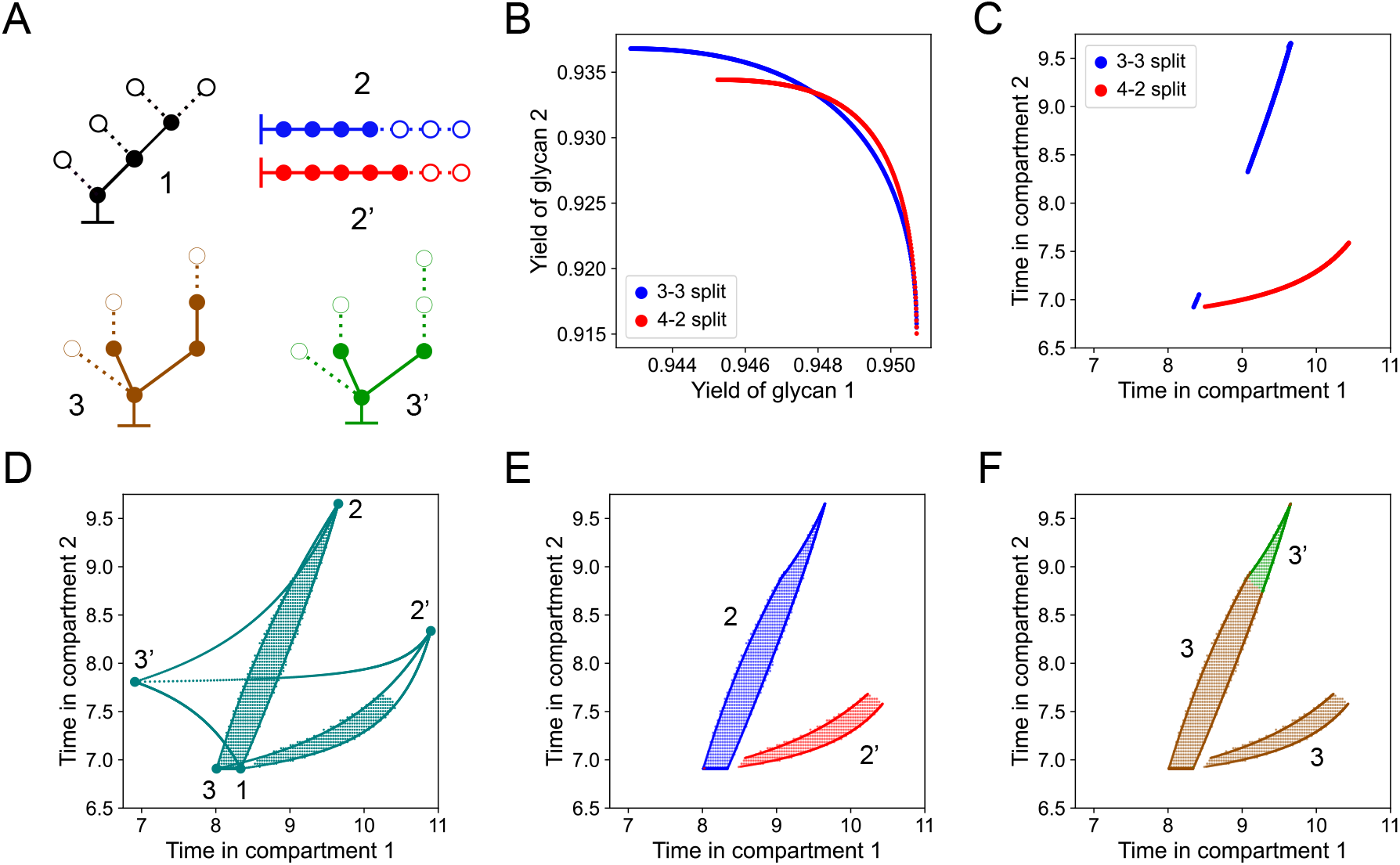
Pareto-optimal splits and residence times for multiple glycan manufacture. **(A)** Pareto-optimal splits of the three trees. Numbers adjacent to the structures and their prime versions (e.g. 2 and 2’) represent different splits of the same tree. The nodes with filled-in colors (along with unbroken lines) represent monomers that will be linked in the first compartment, and the empty nodes with borders (and dotted lines) represent monomers that will be added in the second compartment. The same colors are used to indicate splits in subsequent plots. **(B)** Yields of trees 1 and 2 obtained by maximizing different convex weightings of trees (Appendix B), using split 2 and 2’ (blue and red curves respectively). Tree 1 has only one optimal split. For this and all subsequent plots, *q* = 0.001. **(C)** The Pareto set corresponding to (B). Colors indicate the splits that generated those solutions. **(D)** The labeled points denote optimal residence times for each split. Each thick curve going from one labeled point to another represents Pareto-optimal solutions assuming only those two splits are allowed (and disregarding the third glycan). The dotted region denotes the Pareto-optimal solutions for the three glycans under study. **(E)** and **(F)** Optimal splits of the second and third glycan respectively corresponding to the Pareto set in (D).

### Pareto-optimal solutions for two-glycan manufacture

For two glycans, the Pareto-optimal solutions lie on one-dimensional sets composed of segments associated with distinct enzyme splits. The corresponding Pareto front consists of crossover points, where the optimal enzyme split for glycans switch for a small change in yields. This implies that small shifts in the cell’s relative value of the glycans can change the Pareto-optimal enzyme split.

To see this, consider the Pareto set of trees 1 and 2 shown in Fig. 4A. Candidate solutions were be obtained by fixing the split of both trees and finding the optimal residence times for a range of convex weightings of their log-yields (Appendix B, [43]). This was enumerated for all possible splits of both trees. Fig. 4B shows the yields of both trees under a range of convex weightings of their yields. The blue and red colors denote the different splits of the second tree that are used when calculating the yields (the first tree has one optimal split). Some segments of the blue curve Pareto-dominate segments of the red curve and vice versa [33]. The crossover point represents the point at which the Pareto-optimal split of second tree changes, as the yield of the first glycan is varied. To obtain Pareto-optimal solutions with a higher yield of the second glycan than at crossover point, split 2 (blue) must be used. If the yield of the first glycan is to be higher than at crossover point, then split 2’ (red) must be used. The Pareto set is shown in Fig. 4C.

### Pareto-optimal solutions for three or more glycan manufacture

For three glycans, the Pareto set generalizes to a two-dimensional region in residence-time space. Some such regions are partitioned by internal boundaries where the optimal split of a glycan changes. This implies that small changes in kinetics (and hence in effective residence times) can change the Pareto-optimal enzyme split.

To see this, we first sampled the space of Pareto-optimal solutions for the three glycans in Fig. 4A by exhaustive enumeration. We evaluated the yields for all the splits of the three glycans, across a grid of (*t*_1_, *t*_2_) values. It is sufficient to sample timepoints between the minimum and maximum residence times across all splits of the glycans (Appendix C). At each (*t*_1_, *t*_2_), we obtained the maximum yield across splits for the three glycans. We retained only those splits which produced maximum yield for at least one (*t*_1_, *t*_2_) pair; this substantially reduces further computation needed to find the Pareto-optimal solutions (Appendix C). We also considered candidate solutions generated by pairwise convex combinations of the retained splits [28]: after obtaining the maximizer for a convex weighting, we computed the maximum yield attainable for the third glycan at that maximizer. We then checked for non-dominated solutions across the union of these sets of solutions.

Fig. 4D shows the Pareto-optimal residence times for manufacturing the three glycans. The region is enclosed by boundaries corresponding to candidate solutions generated by pairwise combinations of the retained splits (Fig. 4D, thick teal curves). Not all points inside the boundary specified by the curves are Pareto-optimal, as some points might be Pareto-dominated by other solutions not within the same boundary. Figs. 4E, F show the optimal splits of second and third glycans and residence times that form the Pareto set (there is only one optimal split for glycan 1). Fig. 4F shows that optimal enzyme split might change within the same contiguous region of optimal residence times.

In the three-dimensional yield space, each combination of splits defines a yield manifold. Maximizing different convex weightings of log-yields under those splits gives corresponding optimal residence times. The manifold is obtained by calculating the yields of the three glycans at those residence times. Two such manifolds may intersect along a curve. The intersection curve represents points at which the optimal split changes to achieve differing Pareto-optimal yields; it plays a role similar to the crossover point in two-glycan manufacture (Fig. 4B). If the Pareto-optimal residence times corresponding to two sets of splits happen to be in adjacent sub-regions, the optimal split changes within the same continuous region of the Pareto set (Fig. 4F). The crossover curves where the optimal split changes within a region can be specified analytically and the boundary can be computed numerically, although here we do it by enumeration.

## Discussion

The non-templated, tree-like growth of glycans raises interesting questions about the efficacy of its manufacturing process, such as how enzyme localization and residence time constraints limit attainable yield. Our abstract kinetic formulation addresses this question in a way that is independent of system-specific parameter choices, and is equally applicable to both N- and O-glycans. A Pareto optimality-based perspective allows us to make general statements without invoking a specific objective or fitness function [27], making our conclusions more robust.

In our framework, reconciling the intrinsic trade-offs of glycan biosynthesis corresponds to operating on a Pareto front; we therefore propose that the Golgi might balance competing glycan outputs by operating near Pareto-optimal regimes. The Pareto front is defined by a small number of discrete enzyme splits, even though the number of possible splits is quite large. In engineered compartmentalized reactors, this translates to a practical workflow in which computation narrows the experimental design to a handful of candidate enzyme localizations and predicts residence-time regimes where switching between them becomes beneficial. In case of three glycan manufacture, we conjecture that the Pareto-optimal solutions will be contained in the boundaries obtained by assuming that only two splits are allowed (Fig. 4D).

Notably, small changes in the residence times or relative glycan yields can lead to different optimal splits (Fig. 4B,F). This behaviour is analogous to first-order phase transitions reported in other Pareto-optimized systems [44]; e.g. in three glycan manufacture, when the residence time of second compartment is treated as an externally tuned parameter (with the remaining decision variables re-optimized), the optimal enzyme partition switches, similar to an order-parameter jump. Although the observation also indicates that yields of different splits are comparable, even modest increases in yields may confer a survival advantage as glycan biosynthesis is scaled across a large number of cells. In other contexts where there is a crossover, hedging between two configurations provides higher average objective than either of the configurations alone [33]. We have not explored this possibility of bet hedging in the context of glycan manufacture.

Our model makes three predictions that can be probed in vitro using immobilized enzyme cascades [45] or droplet-based microfluidic platforms [46], originally developed for targeted glycosylation. In these systems, each reactor module or incubation chamber can be viewed as a Golgi compartment, with reaction or chamber time representing residence time and the allocation of enzymes in early and late modules representing an effective enzyme split. First, for a fixed enzyme split, the yield of a given glycan is predicted to be unimodal as a function of compartment residence times (Fig. 2C). This can be tested by sampling a range of effective residence times – for example by changing reaction time in immobilized cascades or adjusting flow rates and the number of incubation chambers in microfluidics – and measuring yields using established glycomics workflows [45, 46]. Second, when multiple glycans are produced simultaneously, the model predicts trade-offs in their yields under different residence times (Fig. 4B). The absence of a non-trivial front (for example, a single configuration that dominates all others) would falsify this prediction. Third, within the Pareto front, small changes in residence times are predicted to change the enzyme split that optimizes the joint yields (Fig. 4F). Our kinetic framework can be applied to identify a small number of candidate enzyme splits in the experimental setup whose predicted yields are similar and have a crossover (Fig. 4B). The residence times can be varied in a small neighborhood around the model-predicted crossover to test for such switching. Performing analogous tests in biological Golgi stacks is more challenging because compartment-specific residence times remain difficult to measure in vivo.

We have considered only partitions of enzymes as candidate splits for tractability. Redundant localization of enzymes will increase the rate of undesirable reactions and its net effect on yield will be context-dependent; a systematic exploration of this possibility is left for subsequent studies. We have assumed for simplicity that enzyme splits can be set independently for each glycan, when in fact enzymes executing desirable reactions can be shared across different glycans [15, 21]. This aspect becomes relevant while considering other models of Golgi trafficking [14], where cargo can have different residence times in the Golgi [47]. The presence of common enzymes involved in the manufacture of different glycans will again create trade-offs. In summary, regardless of the underlying model of Golgi trafficking used, some trade-offs must arise. Adapting our model to these scenarios will enable direct comparison with existing glycan distribution and residence time data [47]. Another interesting framing would be to impose a global time budget for Golgi transit in our model and ask how close observed enzyme distributions lie to a speed–error frontier [48].

Critical quality attributes for therapeutic antibodies specify acceptable glycan profiles and help evaluate their safety and efficacy [49]. Given a glycosylation reaction network and the appropriate parameter values, yields for glycans of interest can be computed [50]. This can be used to generate Pareto-optimal solutions for the glycosylation reaction network under the required constraints. Studying the Pareto set and Pareto front will provide insights into the sensitivity of yields and robustness of the optimizer to parameter changes (Fig. 3A, Fig. 4F). Thus, a Pareto optimality-based framework for glycan manufacture has broad applicability, ranging from addressing fundamental questions about how cells resolve biosynthetic trade-offs, to guiding the design of efficient glycosylation processes for next-generation biotherapeutics.

## Code availability

Calculations were performed using SageMath software [51] and libraries NumPy (version 1.26.4) [52], SciPy (version 1.16.2) [53], and pandas (version 2.2.3) [54]. Plots were generated using matplotlib version 3.8.0 [55]. A Jupyter Notebook generating the plots can be found at: https://gitlab.com/aashishsatya/pareto-optimal-glycan-synthesis.

## Author Contributions

MT conceived the project. AS carried out the analysis. AS wrote the paper.

## Acknowledgments

AS would like to thank Saptarshi Dasgupta for constructive feedback on the manuscript. The authors would like to thank Nandita Chaturvedi, Sachit Daniel, and Shaon Chakraborty for useful discussions.

MT acknowledges support from the Simons Foundation (287975). AS would like to thank Chennai Mathematical Institute for funding.

## Appendix

### A. Finding optimal splits and residence times for single glycan manufacture

*The yield objective is a log-concave function of residence times:* A positive function *f* : ℝ → ℝ_+_ is log-concave if the logarithm of the function is concave, i.e. log *f* (*θx* + (1 − *θ*)*y*) ≥ *θ* log *f* (*x*) +(1 − *θ*) log *f* (*y*) for 0 ≤ *θ* ≤ 1; *x, y* ∈ ℝ [42]. A differentiable function *f* is concave in an interval if its first derivative is monotonically decreasing in that interval. From the definition of log-concavity, it follows that if *f* can be decomposed into a product of log-concave functions, then *f* is log-concave.

A function *f* (*x*_1_, …, *x*_*N*_) is said to be pointwise log-concave if for all fixed *x*_1_, …, *x*_*i*−1_, *x*_*i*+1_, …, *x*_*N*_, the function is log-concave in *x*_*i*_, ∀*i*. We prove that the yield function *yield*( 𝒮, 𝒯) for a glycan, with fixed 𝒮, is point-wise log-concave. Since the yield function comprises products of yields corresponding to splits in each compartment, it suffices to prove that the expression for yield in a single compartment is a log-concave function of the residence time for that compartment. We do this by induction.

#### Base case

The yield corresponding to a single reaction is (1 − *e*^−*t*^) ·*e*^−*qt*^ for some *q* (*t* is the residence time). Both the terms can be shown to be log-concave by taking their respective logarithms and seeing that the second derivative with respect to time is non-negative. Therefore their product is also log-concave.

#### Inductive case

A split can have several independent components, and the final yield is calculated as the product of yields of individual components (multiplied by the undesirable reaction term, which is log-concave). We show that yield of each component is a log-concave function of the residence time.

Let us assume that the yield of a sub-tree *B*_*i*_ attached to a root node is log-concave w.r.t. the residence time *t*, i.e. *yield-formation*(*B*_*i*_; *t*) is log-concave. The overall yield of the branch containing *B*_*i*_ is calculated as

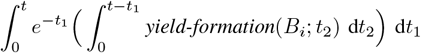

Write 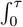 as *Y* (*τ*). The integral of a log-concave function is known to be log-concave [42], thus *Y* (*τ*) is log-concave in *τ*. Then the above expression represents the convolution of two log-concave functions *F* (*τ*) = *e*^−*τ*^ and *Y* (*τ*), which is also known to be log-concave [42].

## B. Generating Pareto-optimal solutions for multiple glycan manufacture

Several methods exist in literature to generate Pareto-optimal solutions in case of concave objectives. If there are multiple concave objectives *f*_1_(***x***), …, *f*_*K*_(***x***); *f*_*i*_(***x***) : ℝ^*n*^→ ℝ for some *n*, then it is known that solutions of

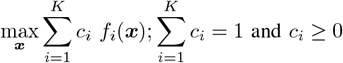

i.e. maximizers of convex combinations of the given concave functions, are Pareto-optimal [57, 58].

We enumerate all possible splits for all glycans (fixing which makes the yield log-concave functions of residence times; see Appendix A) and use exhaustive search to pick the Pareto-optimal solutions. For generating candidates solutions for two-glycan manufacture (Figs. 4B, C), we generated 401 weights uniformly between 0 and 1.

## C Generating Pareto-optimal solutions by exhaustive enumeration

There were 22, 7, and 24 possible splits for each of the trees in Fig. 4A, including splits where the entire tree is built in one compartment. In case of three glycan manufacture, there were 22 × 7 × 24 = 3, 696 possible combinations of splits. To bring down the combination of splits to consider, we evaluated the yields for all the splits of the three glycans, across a grid of (*t*_1_, *t*_2_) values. It suffices to sample timepoints between the minimum and maximum residence times across all splits of the glycans, since yields outside these bounds will be Pareto-dominated:

### Claim

Let 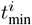 and 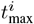 denote the minimum and maximum optimal residence time in a compartment *i* across all splits of *K* glycans to be manufactured. Then 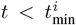 and 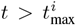 cannot be a Pareto-optimal residence time for that compartment.

**Proof:** It suffices to prove the claim for a single compartment. Consider the log-yields 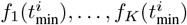 obtained in that compartment for each of the *K* glycans at 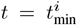. Each of the *K* glycans has optimal residence time greater than or equal to 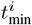. Consider a 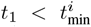. By the concavity of the log-yields, for the *j*^th^ glycan, 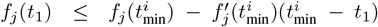 where 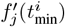 represents the derivative of *f*_*j*_ at 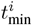 (*f*_*j*_ is differentiable by the differentiability of log function, and the fundamental theorem of calculus). The term 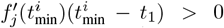 for all the glycans whose optima is not at 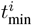. Thus 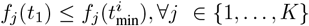, ∀*j* ∈ {1, …, *K*}, and there will exist one or more *j* for which the inequality is strict because of the positive term. Hence, the yields attained at *t*_1_ are Pareto-dominated by those at 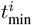, and *t*_1_ will not be Pareto-optimal. An analogous argument can be made for 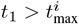.

We sampled a uniformly spaced grid of 251 × 251 values between [6, 14] × [6, 14]. We then obtained the maximum yields for the three glycans at each (*t*_1_, *t*_2_) pair. We retained only those splits which produced maximum yield for at least one (*t*_1_, *t*_2_) pair. This left 7, 5 and 8 splits respectively for the three trees. Among the 251 × 251 = 63, 001 values, the yields at 1, 896 timepoints were non-dominated. We considered 10, 001 weights uniformly spaced between 0 and 1 while considering pairwise combinations of the retained splits (this was needed to filter out any spurious Pareto-optimal solutions outside the boundaries formed by pairwise combinations of splits). The 10, 001 candidates for each 7 × 5 × 8 = 280 combinations along with the 1, 896 solutions above were screened for non-dominated yields, and the Pareto set was plotted in Fig. 4D.

## References

1. Adibekian, A. et al. Comparative bioinformatics analysis of the mammalian and bacterial glycomes. Chemical Science 2, 337–344 (2011).

2. Wetzel, L. A. et al. Predicted glycosyltransferases promote development and prevent spurious cell clumping in the choanoflagellate S. rosetta. eLife 7 (2018).

3. Moremen, K. W., Tiemeyer, M. & Nairn, A. V. Vertebrate protein glycosylation: diversity, synthesis and function. Nature Reviews Molecular Cell Biology 13, 448–462 (2012).

4. Varki, A. Biological roles of glycans. Glycobiology 27, 3–49 (2016).

5. Zhao, X., Chen, H. & Wang, H. Glycans of SARSCoV-2 Spike Protein in Virus Infection and Antibody Production. Frontiers in Molecular Biosciences 8. ISSN: 2296-889X (2021).

6. Lloyd, K. O., Burchell, J., Kudryashov, V., Yin, B. W. T. & Taylor-Papadimitriou, J. Comparison of O-Linked Carbohydrate Chains in MUC-1 Mucin from Normal Breast Epithelial Cell Lines and Breast Carcinoma Cell Lines. Journal of Biological Chemistry 271, 33325– 33334 (1996).

7. Lo-Guidice, J.-M. et al. Sialylation and sulfation of the carbohydrate chains in respiratory mucins from a patient with cystic fibrosis. Journal of Biological Chemistry 269, 18794–18813 (1994).

8. Brockhausen, I. Mucin-type O-glycans in human colon and breast cancer: glycodynamics and functions. EMBO reports 7, 599–604 (2006).

9. Alberts, B. Molecular biology of the cell (Garland Science, 2015).

10. Stanley, P. Golgi glycosylation. Cold Spring Harbor Perspectives in Biology 3 (2011).

11. Glick, B. S. Organization of the Golgi apparatus. Current Opinion in Cell Biology 12, 450–456. ISSN: 09550674 (4 2000).

12. Klumperman, J. Architecture of the Mammalian Golgi. Cold Spring Harbor Perspectives in Biology 3, a005181– a005181. ISSN: 1943-0264 (7 2011).

13. Fisher, P., Spencer, H., Thomas-Oates, J., Wood, A. J. & Ungar, D. Modeling Glycan Processing Reveals Golgi-Enzyme Homeostasis upon Trafficking Defects and Cellular Differentiation. Cell Reports 27, 1231–1243.e6 (2019).

14. Glick, B. S. & Luini, A. Models for Golgi traffic: a critical assessment. Cold Spring Harbor perspectives in biology 3, a005215. ISSN: 1943-0264 (11 2011).

15. Biswas, A. & Thattai, M. Promiscuity and specificity of eukaryotic glycosyltransferases. Biochemical Society Transactions (2020).

16. Aminoff, D., Bruegge, W. F., Bell, W. C., Sarpolis, K. & Williams, R. Role of sialic acid in survival of erythrocytes in the circulation: interaction of neuraminidase-treated and untreated erythrocytes with spleen and liver at the cellular level. Proceedings of the National Academy of Sciences 74, 1521–1524. ISSN: 0027-8424 (4 1977).

17. Varki, A. Sialic acids in human health and disease. Trends in Molecular Medicine 14, 351–360. ISSN: 14714914 (8 2008).

18. Scallon, B. J., Tam, S. H., McCarthy, S. G., Cai, A. N. & Raju, T. S. Higher levels of sialylated Fc glycans in immunoglobulin G molecules can adversely impact functionality. Molecular Immunology 44, 1524–1534. ISSN: 01615890 (7 2007).

19. Schwabl, S. & Teis, D. Protein quality control at the Golgi. Current Opinion in Cell Biology 75, 102074. ISSN: 0955-0674 (2022).

20. Gutierrez, J. M. et al. Genome-scale reconstructions of the mammalian secretory pathway predict metabolic costs and limitations of protein secretion. Nature Communications 11 (2020).

21. Jaiman, A. & Thattai, M. Golgi compartments enable controlled biomolecular assembly using promiscuous enzymes. eLife 9 (2020).

22. Umaña, P. & Bailey, J. E. A mathematical model of Nlinked glycoform biosynthesis. Biotechnology and bioengineering 55, 890–908. ISSN: 0006-3592 (6 1997).

23. Krambeck, F. J. et al. A mathematical model to derive N-glycan structures and cellular enzyme activities from mass spectrometric data. Glycobiology 19, 1163–1175. ISSN: 1460-2423 (11 2009).

24. Spahn, P. N. et al. A Markov chain model for N-linked protein glycosylation – towards a low-parameter tool for model-driven glycoengineering. Metabolic Engineering 33, 52–66 (2016).

25. Krambeck, F. J., Bennun, S. V., Andersen, M. R. & Betenbaugh, M. J. Model-based analysis of Nglycosylation in Chinese hamster ovary cells. PLOS ONE 12, e0175376. ISSN: 1932-6203 (5 2017).

26. Fisher, P., Thomas-Oates, J., Wood, A. J. & Ungar, D. The N-Glycosylation Processing Potential of the Mammalian Golgi Apparatus. Frontiers in Cell and Developmental Biology 7. ISSN: 2296-634X (2019).

27. Yadav, A., Vagne, Q., Sens, P., Iyengar, G. & Rao, M. Glycan processing in the Golgi: optimal information coding and constraints on cisternal number and enzyme specificity. eLife 11 (2022).

28. Miettinen, K. Nonlinear multiobjective optimization (Kluwer Academic Publishers, 1998).

29. Deb, K. Evolutionary Algorithms for Multi-Criterion Optimization in Engineering Design in (1999).

30. Aziz, H., Brandt, F. & Harrenstein, P. Pareto optimality in coalition formation. Games and Economic Behavior 82, 562–581 (2013).

31. Shoval, O. et al. Evolutionary trade-offs, Pareto optimality, and the geometry of phenotype space. Science 336, 1157–1160 (2012).

32. Tendler, A., Mayo, A. & Alon, U. Evolutionary tradeoffs, Pareto optimality and the morphology of ammonite shells. BMC Systems Biology 9, 12. ISSN: 1752-0509 (2015).

33. Levins, R. Theory of fitness in a heterogeneous environment. I. The fitness set and adaptive function. The American Naturalist 96, 361–373 (1962).

34. Jedlicka, P., Bird, A. D. & Cuntz, H. Pareto optimality, economy–effectiveness trade-offs and ion channel degeneracy: Improving population modelling for single neurons. Open Biology 12 (2022).

35. Jefferis, R. Glycosylation as a strategy to improve antibody-based therapeutics. Nature Reviews Drug Discovery 8, 226–234 (2009).

36. Kontoravdi, C. & del Val, I. J. Computational tools for predicting and controlling the glycosylation of biopharmaceuticals. Current Opinion in Chemical Engineering 22, 89–97. ISSN: 22113398 (2018).

37. Zhong, X., D’Antona, A. M., Scarcelli, J. J. & Rouse, J. C. New Opportunities in Glycan Engineering for Therapeutic Proteins. Antibodies 11, 5. ISSN: 2073-4468 (1 2022).

38. Mani, S. & Thattai, M. Stacking the odds for Golgi cisternal maturation. eLife 5 (2016).

39. Varki, A. et al. Essentials of glycobiology 859. ISBN: 9781621824213 (Cold Spring Harbor Laboratory Press, 2022).

40. Pothukuchi, P. et al. GRASP55 regulates intra-Golgi localization of glycosylation enzymes to control glycosphingolipid biosynthesis. The EMBO Journal 40. ISSN: 0261-4189 (20 2021).

41. Neelamegham, S. et al. Updates to the Symbol Nomenclature for Glycans guidelines. Glycobiology 29, 620– 624. ISSN: 1460-2423 (9 2019).

42. Boyd, S. P. & Vandenberghe, L. Convex optimization (Cambridge Univ. Pr., 2011).

43. Emmerich, M. T. M. & Deutz, A. H. A tutorial on multiobjective optimization: fundamentals and evolutionary methods. Natural Computing 17, 585–609. ISSN: 1567-7818 (3 2018).

44. Seoane, L. F. & Solé, R. Phase transitions in Pareto optimal complex networks. Physical Review E 92, 032807. ISSN: 1539-3755 (3 2015).

45. Makrydaki, E. et al. Immobilized enzyme cascade for targeted glycosylation. Nature Chemical Biology 20, 732–741. ISSN: 1552-4450 (6 2024).

46. Isenrich, F. N., Losfeld, M.-E., Aebi, M. & deMello, A. J. Microfluidic mimicry of the Golgi-linked N -glycosylation machinery. Lab on a Chip 25, 1907–1917. ISSN: 1473-0197 (8 2025).

47. Tie, H. C. et al. Quantitative intra-Golgi transport and organization data suggest the stable compartment nature of the Golgi. eLife 13. ISSN: 2050-084X (2025).

48. Chiuchiu, D., Mondal, S. & Pigolotti, S. Pareto optimal fronts of kinetic proofreading. New Journal of Physics 25, 043007. ISSN: 1367-2630 (4 2023).

49. Reusch, D. & Tejada, M. L. Fc glycans of therapeutic antibodies as critical quality attributes. Glycobiology 25, 1325–1334. ISSN: 0959-6658 (12 2015).

50. Flevaris, K., Kotidis, P. & Kontoravdi, C. GlyCompute: towards the automated analysis of protein N-linked glycosylation kinetics via an open-source computational framework. Analytical and Bioanalytical Chemistry 417, 957–972. ISSN: 1618-2642 (5 2025).

51. The Sage Developers. SageMath, the Sage Mathematics Software System (Version 10.5) https://www.sagemath.org (2024).

52. Harris, C. R. et al. Array programming with NumPy. Nature 585, 357–362 (2020).

53. Virtanen, P. et al. SciPy 1.0: Fundamental Algorithms for Scientific Computing in Python. Nature Methods 17, 261–272 (2020).

54. McKinney, W. Data Structures for Statistical Computing in Python in Proceedings of the 9th Python in Science Conference. SciPy 2010 Python in Science ConferenceAustin, Texas (eds van der Walt, S. & Millman, J.) (SciPy, 2010), 56–61.

55. Hunter, J. D. Matplotlib: A 2D graphics environment. Computing in Science & Engineering 9, 90–95 (2007).

56. Mehta, A. Y. & Cummings, R. D. GlycoGlyph: A glycan visualizing, drawing and naming application. Bioinformatics 36, 3613–3614 (2020).

57. Marler, R. T. & Arora, J. S. The weighted sum method for multi-objective optimization: New insights. Structural and Multidisciplinary Optimization 41, 853–862 (2009).

58. Chankong, V. & Haimes, Y. Y. Multiobjective Decision making: Theory and methodology (Dover, 2008).

